# New synthetic-diploid benchmark for accurate variant calling evaluation

**DOI:** 10.1101/223297

**Authors:** Heng Li, Jonathan M Bloom, Yossi Farjoun, Mark Fleharty, Laura Gauthier, Benjamin Neale, Daniel MacArthur

**Author notes:** ^†^To whom correspondence should be addressed., and.

## Abstract

Constructed from the consensus of multiple variant callers based on short-read data, existing benchmark datasets for evaluating variant calling accuracy are biased toward easy regions accessible by known algorithms. We derived a new benchmark dataset from the *de novo* PacBio assemblies of two human cell lines that are homozygous across the whole genome. This benchmark provides a more accurate and less biased estimate of the error rate of small variant calls in a realistic context.

---

Calling genomic sequence variations from resequencing data plays an important role in medical and population genetics, and has become an active research area since the advent of high-throughput sequencing. Many methods have been developed for calling single-nucleotide polymorphisms (SNPs) and short insertions/deletions (INDELs) primarily from short-read data. To measure the accuracy of these methods and ultimately to make accurate variant calls, one typically runs a variant calling pipeline on benchmark datasets where the true variant calls are known. The most widely used benchmark datasets include Genome In A Bottle^1^ (GIAB) and Platinum Genome^2^ (PlatGen) for the human sample NA12878. Both come with a set of high-quality variants and a set of confident regions where non-variant sites are deemed to be identical to the reference genome. These two datasets were constructed from the consensus of multiple short-read variant callers, with consideration of pedigree information or structural variations (SVs) found with long-read technologies. A major concern with GIAB and PlatGen is that the sequencing technologies and the variant calling algorithms used to construct the data sets are the same as the ones used for testing. This strong correlation leads to biases in two subtle ways. First, variant calling with short reads is intrinsically difficult in regions with moderately diverged repeats and segmental duplications. We have to exclude such regions from the confident regions as different variant callers fail to reach consensus there. This biases GIAB and PlatGen toward “easy” genomic regions. In fact, both benchmark datasets conclude competent variant callers make an error every 5 million bases (Figure 1a), while other work suggests we can only achieve an error rate of one per 100–200 thousand bases in wider genomic regions^3,4^, more than an order of magnitude higher. The bias toward easy regions directs the progress in the field to focus on trivial errors while overlooking the major error modes in real applications. Second, as GIAB and PlatGen were constructed using the existing algorithms, they may penalize more advanced algorithms and hamper future method development. For example, FermiKit^5^ achieves high accuracy for INDELs of tens of bases in length that are often missed by other short-read variant callers. Since FermiKit was not one of the callers used in the consensus, the PlatGen dataset is missing many such INDELs and benchmarking against PlatGen will wrongly flag those longer INDELs as false positives (Figure 2a). These caveats suggest we can only comprehensively evaluate the accuracy of short-read variant calling by constructing benchmark datasets with methods orthogonal to and more powerful than short-read sequencing technologies and variant calling algorithms.

**Fig. 1.**
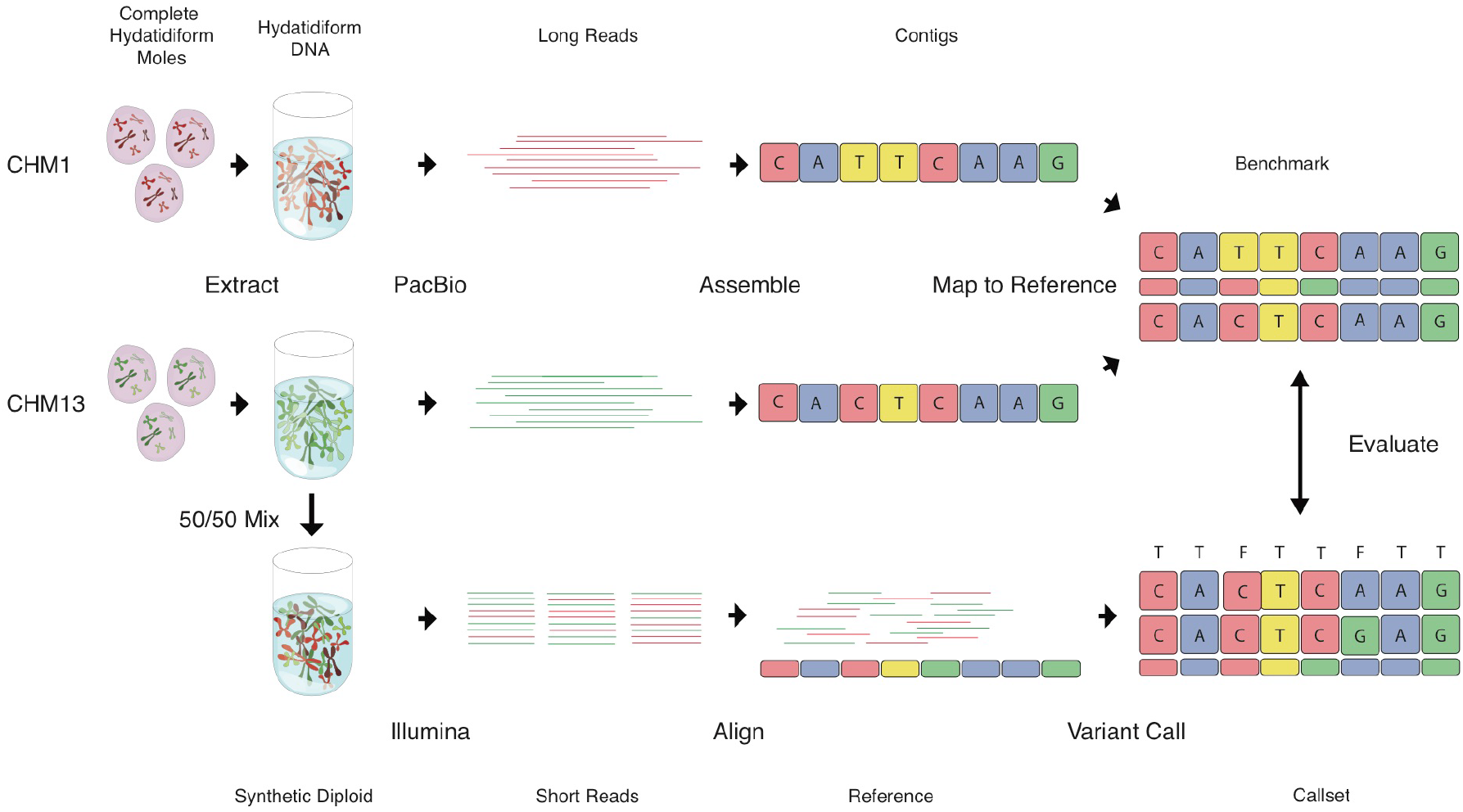
Constructing the Syndip benchmark dataset. CHM1 and CHM13 cell lines were sequenced with PacBio and *de novo* assembled independently. Assembly contigs were aligned to the human reference genome. Differences in the alignment were taken as ‘true’ SNPs and INDELs; regions covered by exactly one contig from each CHM assembly were identified as confident regions where true variants can be called to high accuracy. For the evaluation of diploid variant calling with short reads, equal quantities of DNA from the two cell lines were experimentally mixed. A PCR-free library was constructed from the mix and sequenced to ~40-fold coverage with 151bp paired-end reads. Variants called from the short reads were compared to the PacBio variants as truth to measure variant caller accuracy.

It may be tempting to construct a new benchmark dataset from a whole genome assembly based on PacBio data^6^. However, recent work shows that while PacBio assembly is accurate at the base-pair level for haploid genomes^7^, it is not accurate enough to confidently call heterozygotes in diploid mammalian genomes^8,9^. To derive a comprehensive truth dataset, we turned to the *de novo* PacBio assemblies of two complete hydatidiform mole (CHM) cell lines^10,11^. CHM cell lines are almost completely homozygous across the whole genome. This homozygosity enables accurate PacBio consensus sequences of each cell line and helps to reveal false positives that are caused by copy number variations (CNVs) and manifest as heterozygous SNP and/or INDEL calls on one CHM genome.

Using the “reference genome” of each cell line, we combined the two homozygous calls at each locus into a synthetic diploid call, resulting in the new phased benchmark dataset: Syndip (synthetic diploid; Figure 1). We excluded 1bp INDELs as the PacBio consensus error rate of these INDELs is high, especially around poly-C homopolymer runs. We also excluded poly-A runs ≥10bp in length for a similar reason. In the end, we generated 3.53 million SNPs and 0.38 million 2–50bp INDELs, in 2.70 gigabases (Gbp) of confident regions, covering 95.4% of the autosomes and X chromosome of GRCh37.

In order to compare Syndip with existing benchmarks and re-evaluate popular short-read variant callers, we evenly mixed DNA from the two CHM cell lines and sequenced the mix with Illumina HiSeq X Ten (Figure 1). By counting supporting reads at heterozygous SNPs after variant calling, we estimated 50.7% of DNA in the mixture comes from one cell line and 49.3% from the other, concluding that the mixture is a good representative of a naturally diploid sample. We mapped the reads from the synthetic-diploid samples to the human genome with BWA-MEM-0.7.15^12^, Bowtie2-2.2.2^13^, minimap2-2.5^14^ and SNAP-0.15.7^15^, and called variants on the synthetic-diploid samples with FermiKit-0.1.13^5^, FreeBayes-1.0.2^16^, Platypus-0.8.1^17^, Samtools-1.3^18^ and GATK-3.5^19^, including the HaplotypeCaller (HC) and UnifiedGenotyper (UG) algorithms. We included multiple variant callers to avoid overemphasizing caller-specific effects. We optionally filtered the initial variant calls with the following rules: 1) variant quality ≥ 30; 2) read depth at the variant is below 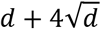 with *d* being the average read depth; 3) the fraction of reads supporting the variant allele ≥ 30% of total read depth at the variant (25% for FermiKit); 4) Fisher strand p-value ≥0.001; 5) the variant allele is supported by at least one read from each strand.

To avoid the complication of different variant representations^1^ and to restrict the types of variants considered in evaluation, we implemented a distance-based approach to measuring the variant accuracy. More precisely, for two variant call sets *A* and *B*, a call in *A* is said to be *found* in *B* if there is a call in *B* that is within 10bp to either side of the call in A. Given a truth and a test callset, a *true positive* (TP) is a true variant also found in the test call set; a *false negative* (FN) is a true variant not found in the test call set; a *false positive* (FP) is a test variant call not found in the truth call set. We define %FNR=100×FN/(TP+FN) and FPPM=10^6^×FP/L, where *L* is the total length of confident regions. Variants not overlapping confident regions, 1bp INDELs and >50bp INDELs are not counted as TP, FN or FP. We took FPPM as a metric instead of the more widely used metric “precision” [=TP/(TP+FN)], because FPPM does not depend on the rate of variation and is thus comparable across datasets of different populations or species.

Figure 2 shows the results of evaluating variant calling pipelines with various benchmarks and conditions. Figure 2a reveals that the FPPM of SNPs estimated from Syndip is often 5–10 times higher than FPPM estimated from GIAB or PlatGen. Looking into the Syndip FP SNPs, we found most of them are located in CNVs that are evident in PacBio data in the context of long flanking regions, but look dubious in short-read data alone. GIAB-3.3.1 and PlatGen-1.0 often exclude these false positives from the truth variant set based on the pedigree information or orthogonal data. However, in real applications, we often only have access to Illumina data and thus cannot achieve the accuracy suggested by the two benchmark datasets.

**Fig. 2.**
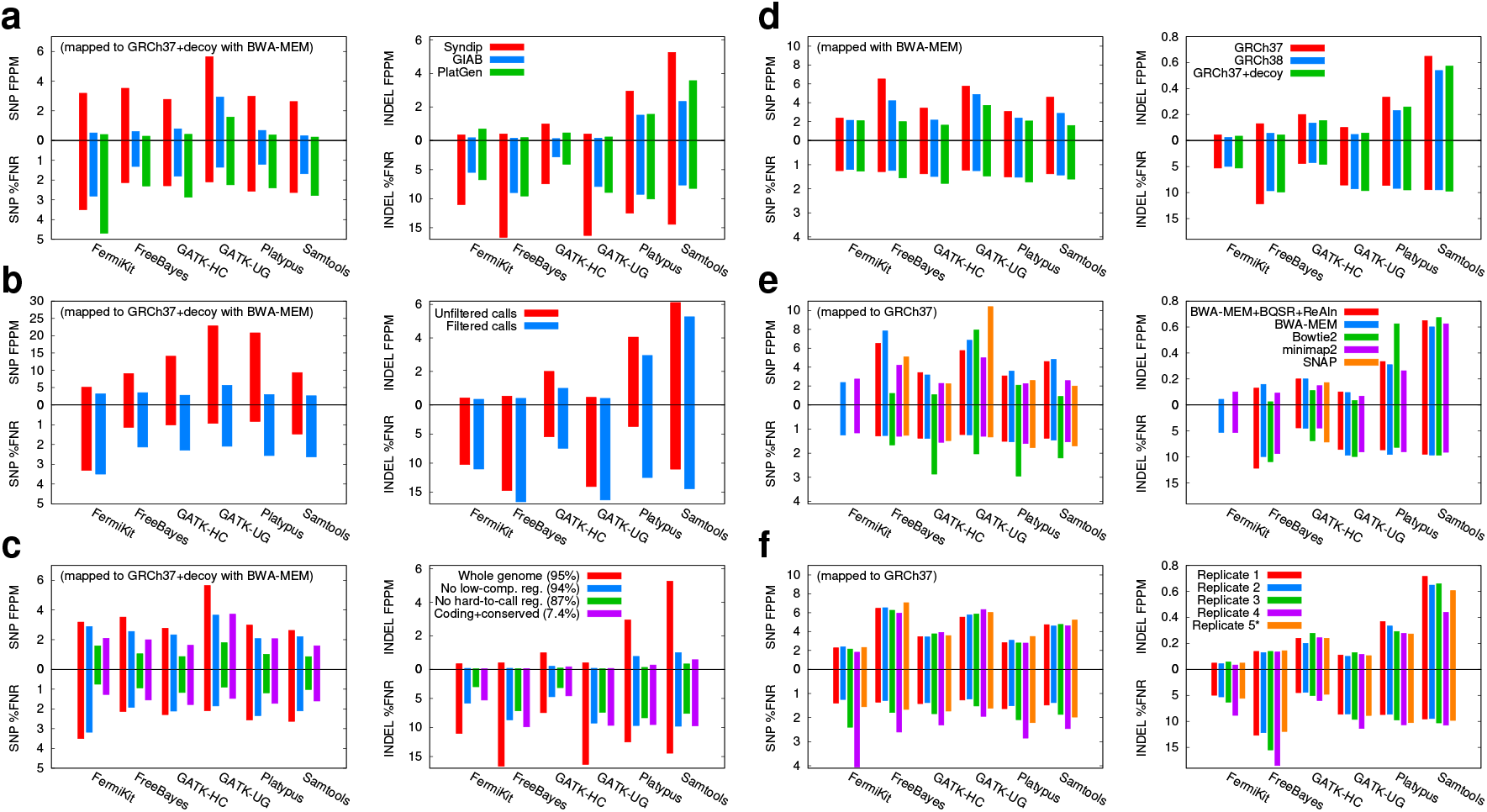
Percent false negative rate (%FNR) and number of false positives per million bases (FPPM) of different variant calling algorithms. (a) Comparison of Syndip, GIAB and PlatGen benchmark data sets on filtered calls. For GIAB and PlatGen, variants were called from the HiSeq X Ten run ‘NA12878_L7_S7’ available from the Illumina BaseSpace. (b) Effect of variant filters. (c) Effect of evaluation regions. Low-complexity regions were identified with the symmetric DUST algorithm^22^ at a score threshold 30. The ‘hard-to-call’ regions include low-complexity regions, regions unmappable with 75bp single-end reads and regions susceptible to common copy number variations^23^. Panels (d)–(f) only show metrics in ‘coding+conserved’ regions. (d) Effect of the human genome reference build. Reference decoy sequences were identified by the 1000 Genomes project. Most of these sequences are real human sequences that are missing from GRCh37. (e) Effect of the mapping algorithms and post-processing. The red data represents variant calls using BWA-MEM for alignment and applied base quality recalibration (BQSR) and INDEL realignment (Realn). INDEL accuracy on SNAP alignment is only shown for GATK-HC because others do not work well with SNAP alignments, which are edit-based. (f) Effect of replication. Replicate 1–4 were sequenced from a library consisting of DNA from both CHM cell lines prior to library construction; Replicate 5^*^ was generated by computationally subsampling and mixing reads sequenced from the two CHM cell lines separately. Replicate 1 is used in panels (a)–(e).

In our evaluation, we used post-filtered variant calls instead of raw calls. For GATK-HC, filtering reduces sensitivity by only 1%, but reduces the number of FPs by five-fold (Figure 2b), reduces the number of coding SNPs absent from the 1000 Genomes Project^20^ by 58%, and reduces the number of loss-of-function (LoF) calls by 30%. We manually inspected 20 filtered LoF calls in IGV^21^ and confirmed that all of them are either false positives or fall outside confident regions; those outside confident regions look spurious as well. False positives are enriched among LoF calls because real LoF mutations are subjected to strong selection but errors are not. For functional analyses, such as in the study of Mendelian diseases, we strongly recommend applying stringent filtering to avoid variant calling artifacts. We note that the popular metric F1-score, which is the average of sensitivity [=TP/(TP+FP)] and precision, is usually higher for unfiltered calls. For example, on GIAB, the F1-score of unfiltered GATK-HC SNP calls is 0.998, higher than that of filtered calls 0.991. The F1 metric may not reflect the accuracy important to clinical applications.

Consistent with our previous finding^3^, most FP INDELs come from low-complexity regions (LCRs), 2.3% of human genome (Figure 2c). While this finding helps to guide our future development, it over-emphasizes a class of INDELs that often have unknown functional implications. To put the evaluation in a more practical context, we compiled a list of potentially functional regions, which consist of coding regions with 20bp flanking regions, regions conserved in vertebrate or mammalian evolution and variants in the ClinVar or GWAScatalog databases with 100bp flanking regions. Only 0.5% of these regions intersect with LCRs. As a result, the FPPM of INDELs in these regions is much lower.

We found that mapping reads to GRCh38 leads to slightly better results than mapping to GRCh37 (Figure 2d), potentially due to the higher quality of the latest build. Although mapping to GRCh37 with decoy sequences further helps to reduce FP calls, this often comes at a minor loss in sensitivity. The choice of read mapping pipelines affects variant calling accuracy more (Figure 2e). Bowtie2 alignment often yields lower FPPM because Bowtie2 intentionally lowers mapping quality of reads with excessive mismatches, which helps to avoid FPs caused by divergent CNVs, but may lead to a bias against regions under balancing selection or reduce sensitivity for species with high heterozygosity. It would be preferable to implement a post-alignment or post-variant filter instead of building the limitation into the mapper. We observed comparable FPPM but varying sensitivity across four biological replicates (Figure 2f). Replicate 4 has the lowest coverage and base quality as well as the lowest variant calling sensitivity. Importantly, replicate 5^*^ in Figure 2f suggests that computationally subsampling and mixing reads sequenced from each CHM cell line separately, which is an easier technical exercise than experimentally mixing DNA to a precise fraction, is adequate for the evaluation of short variant calling.

We have manually inspected FPs and FNs called by each variant caller. GATK-HC performs local re-assembly and consistently achieves the highest INDEL sensitivity (Figure 2). However, it may assemble a spurious haplotype around a long INDEL in a long LCR and make a false INDEL call distant from the truth INDEL. We believe this can be improved with a better assembly algorithm as FermiKit, which also performs assembly, is less affected. FreeBayes is efficient and accurate for SNP calling. However, it does not penalize reads with intermediate mapping quality as much as other variant callers, which may lead to high FPPM in regions affected by CNVs. Platypus and SAMtools also demonstrate good SNP accuracy. Nonetheless, they both suffer from an error mode in which they may call a weakly supported false INDEL that is similar but not identical to a true INDEL ≥10bp away. This affects their FPPM. It is not obvious how to filter such false INDELs without looking at the underlying alignments.

Syndip is a special benchmark dataset that has been constructed from high-quality PacBio assemblies of two independent, homozygous cell lines. It leverages the power of long-read sequencing technologies while avoiding the difficulties in calling heterozygotes from relatively noisy data. Syndip is the first benchmark dataset that does not heavily depend on short-read data and short-read variant callers, and thus more honestly reflects the true accuracy of such variant callers. On the other hand, Syndip also has weakness: the PacBio consensus of homozygous genomes is still associated with a small error rate. Errors in the consensus may incorrectly appear to be false negatives in short-read call sets. We excluded 1bp INDELs and INDELs in long poly-A runs to alleviate this problem but consequently lost the ability to evaluate such INDELs. In addition, due to misassemblies in PacBio data and the difficulties in interpreting SVs, Syndip does not cover the entire genome and is still biased toward relatively easy genomic regions. Better PacBio assembly and long-read based SV calling may further improve the Syndip benchmark dataset.

*Data availability:* Illumina reads from this study were deposited to ENA under accession PRJEB13208. This includes one run for each CHM cell line and four runs for experimental mixtures. Syndip variant calls, confident regions and evaluation script can be downloaded from https://github.com/lh3/CHM-eval.

## Methods

### Identifying misassembled regions on PacBio contigs

We acquired CHM1 and CHM13 *de novo* assemblies (accession GCA_001297185 and GCA_000983455, respectively) from NCBI and downloaded Illumina short reads from SRA (accession SRR2842672 and SRR3099549 for CHM1; SRR2088062 and SRR2088063 for CHM13). We assembled Illumina data with FermiKit, mapped to the corresponding PacBio assemblies and called 60,682 heterozygous substitutions from CHM1 and 187,267 from CHM13. These heterozygous substitutions are often close to each other and have higher-than-average Illumina read depth. They are probably due to misassemblies in the PacBio assembly. We hierarchically clustered heterozygous events as follows: we merged two clusters adjacent on the PacBio assembly if 1) the minimal distance between them is within 10kb and 2) the density of heterozygotes in the merged cluster is at least 1 per 1kb. We identified about 3,000 clusters containing three or more heterozygotes from each PacBio assembly, and softly masked these clustered regions to avoid them complicating downstream variant calling.

### Constructing the truth call set and confident regions

For each CHM PacBio assembly, we split the contigs into 200kb subsequences without overlaps and mapped the split sequences to GRCh37 with BWA-MEM with the ‘-x intractg’ option. To call pseudo-diploid variants from PacBio assemblies, we merged the assembly-to-reference alignments of CHM1 and CHM13. We discarded alignments with mapping quality below 20, dropped aligned segments shorter than 10kb and made an unfiltered call set by calling the alignment differences between each PacBio contig and GRCh37.

We constructed the initial set of confident regions from the same alignment. For each PacBio assembly, we say a region on GRCh37 is *orthologous* to the assembly if 1) the region is covered by one PacBio alignment longer than 10kb with mapping quality at least 20; 2) the region is not covered by another PacBio alignment longer than 1kb, regardless of the mapping quality; 3) the aligned position on the PacBio contig is not in a previously identified misassembled region. The initial set of confident regions is the intersection of GRCh37 regions orthologous to both CHM1 and CHM13. These regions cover 96.2% of GRCh37.

In downstream evaluation, we later noticed that if a small region harbors excessive variant calls, the region tends to be enriched with errors potentially due to misalignments or structural variations. We thus applied another hierarchical clustering to spot clusters of variations. More precisely, in this clustering procedure, we merged two clusters if 1) the minimal distance between two variants is within 250bp and 2) the density of variants in the merged cluster is at least 1 per 50bp. We collected clusters consisting of 10 or more variants and excluded the related regions from the initial confident regions. The final confident regions cover 95.9% of GRCh37, and 95.4% when we also excluded poly-A runs ≥10bp. We applied a similar procedure to both GRCh37 with decoy contigs and GRCh38.

To confirm the quality of the Syndip data set, we manually inspected several hundred discordant calls in IGV^21^. We observed that 10–20% of false positive and false negative INDEL calls made by HaplotypeCaller appear to have strong support from Illumina reads. Most of them are around low-complexity regions (LCRs) and supported by both Illumina reads sequenced by us and Illumina reads downloaded from SRA. We speculate that the PacBio-Illumina differences are enriched with consensus errors in PacBio contigs, though somatic mutations and systematic Illumina errors may also contribute. Regardless, under the assumption of perfect Illumina data, 10–20% discrepancy between Illumina and PacBio evidence would not change our general conclusions or the relative performance between calling methods as PacBio contig errors and somatic mutations are not biased toward a particular calling method.

### Quantification, normalization and mixing of the CHM samples

Initial sample quantification was performed using the Invitrogen Quant-It broad range dsDNA quantification assay kit (Thermo Scientific Catalog: Q33130) with a 1:200 PicoGreen dilution. Following quantification, each sample was normalized to a concentration of 10 ng/μL using a 1X Low TE pH 7.0 solution, then sample concentration was confirmed via PicoGreen. Sample mixing was then performed by combining an equal mass (ng) of each of the two samples (CHM1 & CHM13) needed to obtain enough material for the Whole Genome library preparation (500ng). The samples for creating the 4 libraries were normalized and mixed independently.

### Preparation of libraries & sequencing

For PCR-free whole genomes, library construction was performed using Kapa Biosystems reagents with the following modifications: (1) initial genomic DNA input was reduced from 3μg to 500ng, and (2) custom full-length dual-indexed library adapters at a concentration of 15 uM were utilized. Following sample preparation, libraries were quantified using quantitative PCR (kit purchased from Kapa biosystems) with probes specific to adapter ends in an automated fashion on Agilent’s Bravo liquid handling platform. Based on qPCR quantification, libraries were normalized and pooled on the Hamilton MiniStar liquid handling platform. For HiSeq X Ten, pooled samples were normalized to 2nM and denatured with 0.1N NaOH for a loading concentration of 200 pM. Cluster amplification of denatured templates and paired-end sequencing was then performed according to the manufacturer’s protocol (Illumina) for the HiSeq X Ten, with the following modification: we enabled dual indexing outside of the standard HiSeq control software by altering the sequencing recipe files.

### Calling SNPs and short INDELs from Illumina data

We mapped the Illumina reads to the human genome GRCh37 with the GATK best-practice pipeline, which uses BWA-MEM for mapping and post-processes alignments with BQSR and INDEL realignment. We additionally mapped the reads from one sample with BWA-MEM to various human genome versions without post processing steps. We have also run minimap2, Bowtie2 and SNAP for the same sample. We used the default settings of various mappers, except for tuning the maximal insert size.

We called variants on the mixed synthetic-diploid samples with FermiKit, FreeBayes, Platypus, Samtools and GATK, including the HaplotypeCaller (HC) and UnifiedGenotyper (UG) algorithms and filtered the raw variant calls with the set of rules described in the main text. We have tried GATK’s VQSR model for filtering. However, as the VQSR training set is biased towards variants in regions with unambiguous mapping, VQSR misses many truth variants without perfect averaged mapping quality. Both GATK and Platypus come with a set of hard filters. However, by not filtering on read depth, one of the most effective filters on single-sample WGS calling, these filters lead to a low precision.

The variant calling pipeline and filters have been implemented in https://github.com/lh3/unicall.

